# Identification and classification of the genomes of novel Microviruses in poultry slaughterhouse

**DOI:** 10.1101/2024.01.22.576691

**Authors:** Ke-Ming Xie, Ben-Fu Lin, Peng Zhu, Xin-Yu Sun, Chang Liu, Guang-Feng Liu, Xu-Dong Cao, Jing-Qi Pan, Sui-Ping Qiu, Xiao-Qi Yuan, Meng-Shi Liang, Jing-Zhe Jiang, Li-Hong Yuan

## Abstract

Microviridae is a family of phages with circular ssDNA genomes and they are widely found in various environments and organisms. In this study, Virome techniques were employed to explore potential members of Microviridae in poultry slaughterhouse, leading to the identification of 98 novel and complete microvirus genomes. Using a similarity clustering network classification approach, these viruses were found to belong to at least 6 new subfamilies within Microviridae and 3 higher-level taxonomic units. Analysis of their genomes found that the genome size, GC content and genome structure of these new taxa showed evident regularities, validating the rationality of our classification method. Compared with the 19 families classified by previous researchers for microviruses dataset, our method can divide microviruses into about 45 more detailed clusters, which may serve as a new standard for classifying Microviridae members. Furthermore, addressing the scarcity of host information for microviruses, this study significantly broadened their host range and discovered over 20 possible new hosts, including important pathogenic bacteria such as Helicobacter pylori and Vibrio cholerae, as well as different taxa demonstrated differential host specificity. The findings of this study effectively expand the diversity of the Microviridae, providing new insights for their classification and identification. Additionally, it offers a novel perspective for monitoring and controlling pathogenic microorganisms in poultry slaughterhouse environments.

## 1 Introduction

China is a major player in livestock and poultry farming and consumption. According to the statistics, China’s total meat consumption is nearly 100 million tons, accounting for 27% of the global total. In 2022, the domestic meat production reached 92.27 million tons, with poultry meat contributing 24.43 million tons, constituting 26.5% of the total global meat production(1).

Slaughterhouses play a crucial role as an essential pathway for livestock and poultry meat products to move from farms to consumers’ tables. They are also the key points for the gathering and transmission of pathogenic microorganisms(2). Due to the high density and mobility of poultry when entering the market or slaughterhouses, poultry comes from diverse sources and has varying hygienic conditions, and may carry multiple pathogenic microorganisms(3). The slaughter process is prone to contaminating the environment and the personnel involved. Additionally, the waste generated during poultry slaughter and processing further provides favorable conditions for the proliferation of pathogenic microorganisms(4). The interaction between animals, the environment, and occupational personnel forms a closed-loop microbial transmission chain. Some pathogenic microorganisms can infect occupational personnel through direct contact, while others may have an indirect impact by contaminating the environment. Existing research indicates that the detection rate of certain pathogenic microorganisms, such as Campylobacter and Salmonella in the case of bacteria, avian influenza viruses in the case of viruses, is significantly higher among occupational personnel in comparison to the general population(5–9). Therefore, conducting extensive microbial research at the interface of animals, the environment, and occupational personnel in poultry slaughterhouses is of significant importance.

Bacteriophages, a type of viruses that specifically infect bacteria, are the most abundant life forms on earth(10). It is estimated that there are as many as 10^31^ virus particles on earth(11, 12), representing a vast and largely untapped reservoir of biological resources. In the preliminary research conducted by our research group, we identified a significant number of pathogens from poultry slaughterhouse samples(2), along with a vast amount of novel bacteriophages (unpublished data). On the one hand, the abundant presence of pathogens in slaughterhouses creates favorable conditions for the survival of bacteriophages. Investigating the diversity, types, and hosts of bacteriophages in poultry slaughterhouses can enhance our understanding of the composition, transmission, and the interplay between pathogens and bacteriophages in such a unique environment. On the other hand, in poultry slaughterhouses, occupational personnel are at the core of operations, and exposures to pathogenic bacteria increase the risk of infections of this particular group of population, which is a major public health safety hazard. Therefore, it would be advantageous to fully explore and develop potential functional phage species based on the high diversity of phages in poultry slaughterhouses, we can effectively purify the environment, block the spread of pathogenic bacteria in poultry slaughterhouses to safeguard public health safety.

Members of the Microviridae are one of the most widely distributed single-stranded DNA viruses and their natural hosts include pathogenic bacteria such as Spiroplasma, Chlamydia, and Enterobacteria(13). Despite earlier limited attention to the Microviridae, recent research indicates their significant importance in the virosphere(14). At present, the only subfamilies in Microviridae recognized by ICTV are Bullavirinae and Gokushovirinae(15), which cannot fully reflect the diversity of viruses in this familyWhile more Microviridae subfamilies, such as Alpavirinae(16) and Pichovirinae(14) have recently been proposed, the number of classified groups and host information about Microviridae remain severely limited in the literature. This study takes the unique and biologically significant environment of a poultry slaughterhouse in Guangzhou in Guangdong Province in China and employs a multiple displacement amplification (MDA) method(17). Combined with metagenomics sequencing to obtain environmental virus sequencing data from the poultry slaughterhouse (DSV, Dataset of Slaughterhouse Virome) in Guangzhou. Within this dataset, we discovered a diverse set of novel viruses belonging to the Microviridae. A detailed analysis of 98 nearly complete Microviridae genomes revealed their classification into at least six new subfamilies and three higher-level taxonomic units. Comparative analysis with publicly available viral databases demonstrated the high resolution of our classification. Additionally, over 20 potential hosts for microviruses were identified. This study expands our knowledge of the evolution, diversity, and host range of Microviridae, providing insights into the potential biosecurity and ecological significance of these microviruses in poultry slaughterhouses.

## 2 Materials and Methods

### 2.1 Sample Collection

Atotal of three types of samples were collected from a poultry slaughterhouse in a district of Guangzhou: animals, occupational personnel, and environmental samples. The environmental samples included air, soil, sludge, swabs from transportation vehicles, and swabs from the slaughterhouse workshop. The collection protocols were as follows: (1) Animal Samples: Sterile cotton swabs were inserted into the oral cavity and cloaca of chickens or ducks, rotated three times, and then removed. The swab’s tail was discarded, and the swab was immersed in sterile 0.5% BSA-PBS buffer for preservation. Three chickens or ducks from each of the three spaces (caged area, pre-slaughter area, slaughter area) had their oral and cloacal swabs mixed to form one sample. (2) Occupational Personnel Nasal Swab Samples: To collect nasal swab samples from occupational personnel, a sterile cotton swab was gently inserted into the nasal pharynx of the participating volunteer. After a few seconds, the swab was gently rotated and removed. The swab’s tail was discarded, and the swab was immersed in sterile 0.5% BSA-PBS buffer for preservation. Nasal swab samples from a single person with both nostrils were placed in an individual sample collection tube. Written informed consent was obtained from all participants. (3) Air Samples: BioSamplers KIT (225-9595, SKC, Eighty Four, PA) were installed at approximately 1.5 meters above floor at the ventilation points in the slaughter area, pre-slaughter area, and caged area (3 sampling points in total). Using 0.5% BSA-PBS buffer at a flow rate of 8 mL/h, sampling was conducted for 12 hours per day at 110V. Each day’s collection was considered one air sample, and this process was repeated continuously for 3 days. The collected samples were stored in PBS buffer. (4) Soil Samples: Soil samples were collected using the quincunx sampling method at various spaces(18), including the entrance of the poultry slaughterhouse, the slaughter workshop, and the pre-slaughter caged area. Each sample weighed 5-10g. (5) Sludge Samples: Sludge samples were collected at the four corners of the sewage discharge pool, with approximately 10 mL of sewage collected per sample. (6) Environmental Swab Samples: Sterile cotton swabs were used to collect environmental samples from the slaughter workshop, pre-slaughter area, caged area, and poultry transportation vehicles. Five swab samples were collected from each space or vehicle, discarding the swab tails and placing them in sterile 0.5% BSA-PBS buffer for preservation. After collection, all samples were stored at 4°C, transported to the laboratory in a cooler, and then stored long-term at −80°C in an ultra-low-temperature freezer. This study was approved by the Medical Ethics Committee of the School of Public Health, Sun Yat-sen University (Permit No. [2018] No. 001).

### 2.2 Sample Pool Preparation

In order to analyze the virus content and types in samples from different spaces (i.e. slaughter area, pre-slaughter area, caged area) and different types (i.e. air, animals, sludge.) within the slaughterhouse, we combined samples of the same type collected from the same space to prepare sample pools: (1) Combined oral and cloacal swabs from 20 ducks in each of the three spaces (caged area, pre-slaughter area, slaughter area) to create a pool (total of 60 ducks). Combined oral and cloacal swabs from 30 chickens in each of the three spaces to create a pool (total of 90 chickens). (2) Combined nasal swab samples from 20 frontline slaughterhouse workers into one pool. (3) Combined air samples collected continuously for 3 days from each sampling point, creating one pool per sampling point. (4) Combined soil samples collected from each space (mixed samples with four or more points) into one pool. (5) Combined sludge samples collected from each sewage discharge pool into one pool. (6) Combined swab samples from seven slaughterhouse process points in the workshop into one pool. (7) Combined swab samples collected from three poultry transportation vehicles into one pool. Sample pool information is provided in Supplementary Table S2.

**Table 1.**
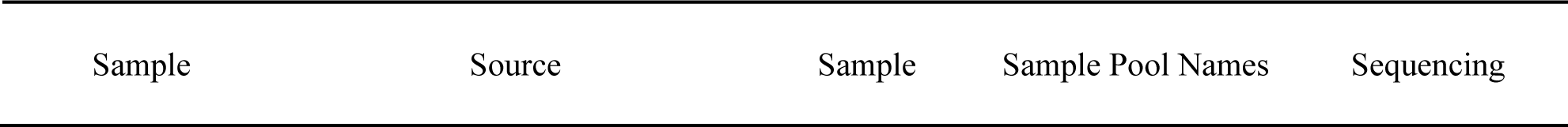
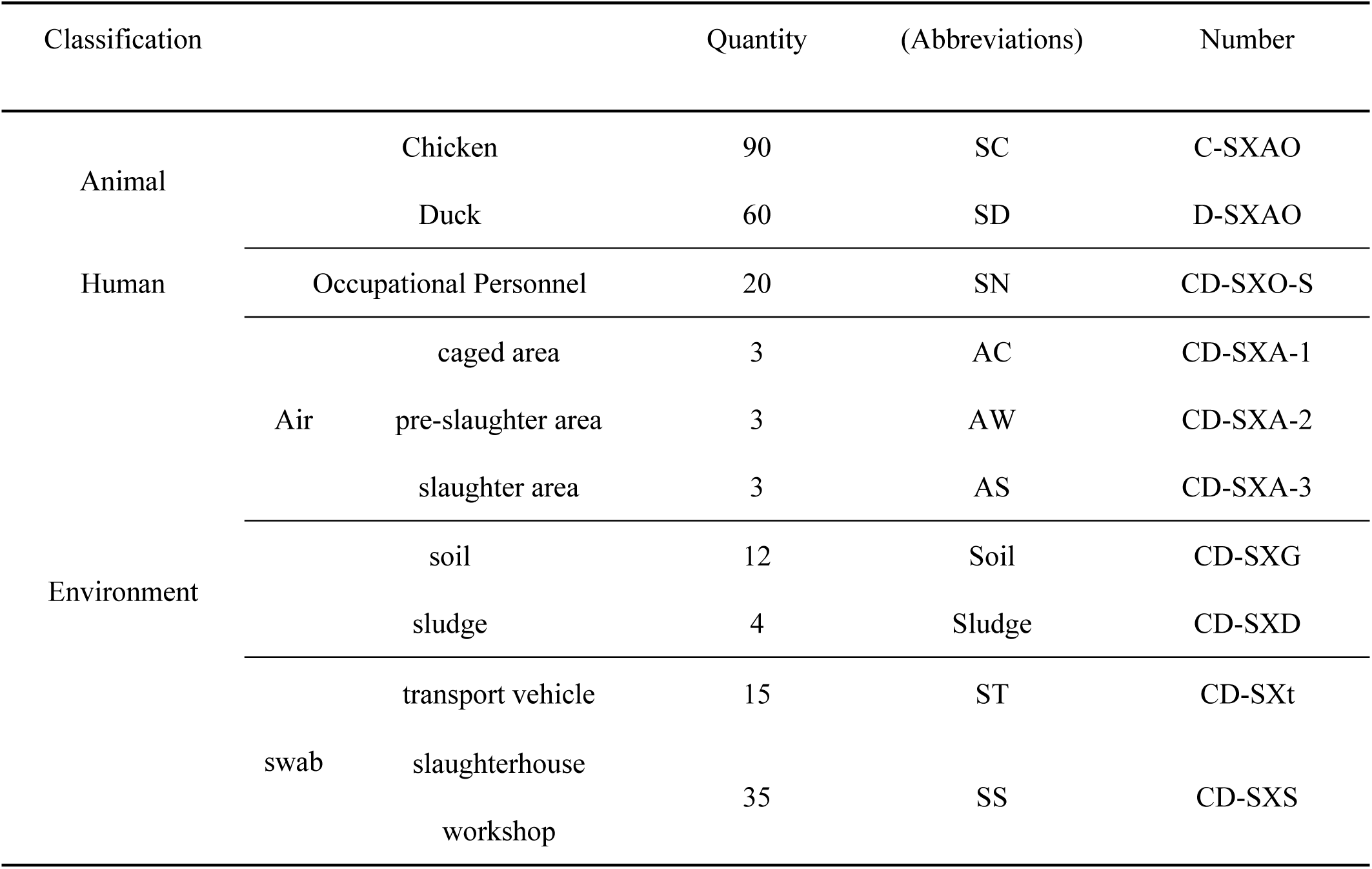
Sample information.

### 2.3 Virus Enrichment, Nucleic Acid Extraction and Amplification

Virus-like particles (VLPs) were enriched separately based on the different properties of the samples. Approximately 0.4g of sludge and soil samples were taken, and each was added to about 2–5 volumes of sterile SB buffer (0.2 M NaCl, 50 mM Tris-HCl, 5 mM CaCl2, 5 mM MgCl2, pH 7.5). For air samples and swab samples, they were directly added to 2–5 volumes of sterile SB buffer and shaken to fully dissolve the virus particles. After three cycles of freeze-thawing, the particles were completely resuspended in 10 times the volume of pre-chilled SB buffer. All samples were centrifuged at 1,000, 3,000, 5,000, 8,000, 10,000, and 12,000 × g for 5 minutes at 4°C using a Sigma 3K30 centrifuge (Sigma Laborzentrifugen GmbH, Germany), and the supernatant was collected. Subsequently, the supernatant was filtered through 0.22 μm Millipore filters (Burlington, MA) to further remove any cell debris and organelles. The filtrate was transferred to 28% sucrose solution and ultra centrifuged at 300,000 × g for 2 hours in a Himac CP 100WX ultracentrifuge (Hitachi, Tokyo, Japan). The supernatant was discarded, and the pellet was re-suspended in 720 μl of water, 90 μl of 10 × DNase I Buffer, and 90 μl of DNase I (1 U/μl) (TAKARA, Japan). The suspension was thoroughly re-suspended, incubated at 37°C with shaking for 60 minutes, stored overnight at 4°C, and then transferred to a 2 ml centrifuge tube.

Total nucleic acids were extracted using the HP Virus DNA/RNA Kit (R6873; Omega Bio-Tek, Norcross, USA), and carrier RNA was not used during the process to avoid potential interference with sequencing results. The concentration of RNA was quantified using the Qubit™ dsDNA HS Assay Kit (Q32851) and Qubit™ RNA HS Assay Kit (Q32855) (Thermo Fisher Scientific, Waltham, USA).

Virome research heavily relies on amplification, as the viral biomass in natural samples is often very low. Due to variations in most amplification methods, quantitative studies of viral data present challenges at present(19, 20). In the current study, uniform genome amplification (WGA) and transcriptome amplification (WTA) were performed using the repi-g Cell WGA and WTA Kit (150052, Qiagen, Hilden, Germany), based on the multiple displacement amplification (MDA) method(17, 21–23).

### 2.4 Library Construction and Sequencing

The amplified DNA was quantified using gel electrophoresis and Nanodrop 2000 spectrophotometer (Thermo Fisher Scientific, Waltham, MA). Ultrasonic random shearing (Covaris M220) was performed to generate fragments with lengths ≤800 bp. Fragment ends were repaired using T4 DNA Polymerase (M4211, Promega, Madison, Wisconsin), Klenow DNA Polymerase (KP810250, Epicentre, Madison, Wisconsin), and T4 Polynucleotide Kinase (EK0031, Thermo Fisher Scientific, Waltham, MA). Fragments in the range of 300-800 bp were collected after electrophoresis. After amplification, the libraries were pooled, and paired-end sequencing of 150 bp, 250 bp, or 300 bp was performed on the Novaseq 6000, HiSeq X ten, and Miseq platforms (Illumina, San Diego, California)(24–26).

### 2.5 Sequence Filtering

All samples underwent metavirome sequencing, yielding approximately 700 million raw sequence reads. The sequencing data were subjected to quality control and removal of low-quality and adapter sequences using Fastp (version 0.20.0) (27). The reads were then assembled into contigs using Megahit (version 1.2.9) (28, 29). The contigs were aligned and annotated against the NCBI non-redundant protein database using Diamond (version 0.9.24.125) (30). Subsequently, Megan6was employed for further classification of the annotated results(31). A total of 98 viral sequences (Contig ID are shown in Table S1) were identified as complete genomes and annotated as belonging to the Microviridae for further in-depth analysis.

### 2.6 Open Reading Frame (ORF) Prediction and Alignment

Cenote-Taker2 was used to predict open reading frames (ORFs) in the 98 viral genomes(32). The major capsid protein or capsid protein sequence (major capsid protein is preferred if available, otherwise capsid protein is chosen. These proteins are collectively referred to as “Cap”) was selected from the predicted results of each viral sequence. NCBI BLASTP(33, 34) was used to compare ORF sequences with the NR database, with an Expect threshold (e-value) set to 10^-5^. For each ORF in the alignment results, the top ten protein sequences with their complete genomic sequences were downloaded based on identity. Duplicate sequences were removed from all downloaded sequences. SnapGene (www.snapgene.com, version 4.3.6) was utilized to open the Cenote-Taker2 output file for visualizing the genomic structure.

### 2.7 Clustering Analysis Based on Sequence Similarity

Cap sequences predicted for DSV microvirus were collected, along with top 10 ranked Cap sequences from the aforementioned BLASTP results, and introduced 20 Cap sequences from microviruses that have been definitively classified by the ICTV. A total of 577 sequences were aligned with each other using DIAMOND (version 0.9.14.115) to build a matrix of sequence similarities. A clustering network graph was constructed based on alignment scores using Gephi(35) (version 0.9.7). The nodes were colored on different sequence sources, hosts, or virus classification results. Furthermore, the Cap sequences used by Paul et al. (36) were integrated with the above data. The same method was employed to construct a similarity clustering network graph and color it, aiming to compare the network clustering resolution of our research method with that of Paul et al.

### 2.8 Host prediction

All complete genome sequences included in the analysis in section 2.7 were subjected to host prediction using hostG(37) (output results taking genus, genus_score > 0.7) and cherry(38) (output results taking Top_1_label, Score_1 > 0.7), analyzing the relationship between these viruses and their hosts, as well as the proportion of these hosts.

### 2.9 Phylogenetic Tree Based on Cap Sequences

Cap is a conserved gene of microviruses(39) with approximately 500 amino acids in length, and is commonly used as a phylogenetic marker for the classification of evolutionary branches or subfamilies within the Microviridae. Multiple sequence alignment was performed using MAFFT(40) ambiguous regions were removed using TrimAl(41), and a maximum likelihood phylogenetic tree based on Cap sequences was constructed using IQtree(42) (version 2.1.4). ModelFinder(43) was set to MFP (for ModelFinder Plus), and 1000 ultrafast bootstrap replicates were used. The tree was visualized using iTOL(44) (version 6.5.2) (https://itol.embl.de).

### 2.10 Principles of Classification and Naming of Viral Sequences

According to the clustering in Figure 1 and cherry host prediction results, DSV-related viral sequences are named respectively. Taking cluster_1 as an example, if a sequence has host prediction results, it is named based on the host, such as the contig sequences CD-SXS-WGA-1-k141_397009 and CD-SXD-WGA-1-k141_230904 are named Bdellovibrio microvirus C1_1 and Bdellovibrio microvirus C1_2, respectively. Similarly, CD-SXG-WGA-1-k141_33139 and CD-SXG-WGA-1-k141_32996 are named Escherichia microvirus C1_1 and Escherichia microvirus C1_2. If the sequence has no host prediction results, contig sequences like CD-SXD-SXG-WGA-all--k141_113185 and CD-SXD-SXG-WGA-all--k141_328845 are named DSV microvirus C1_1 and DSV microvirus C1_2, and so forth. The original sequence ID and their corresponding names are listed in Table S1.

**Figure 1.**
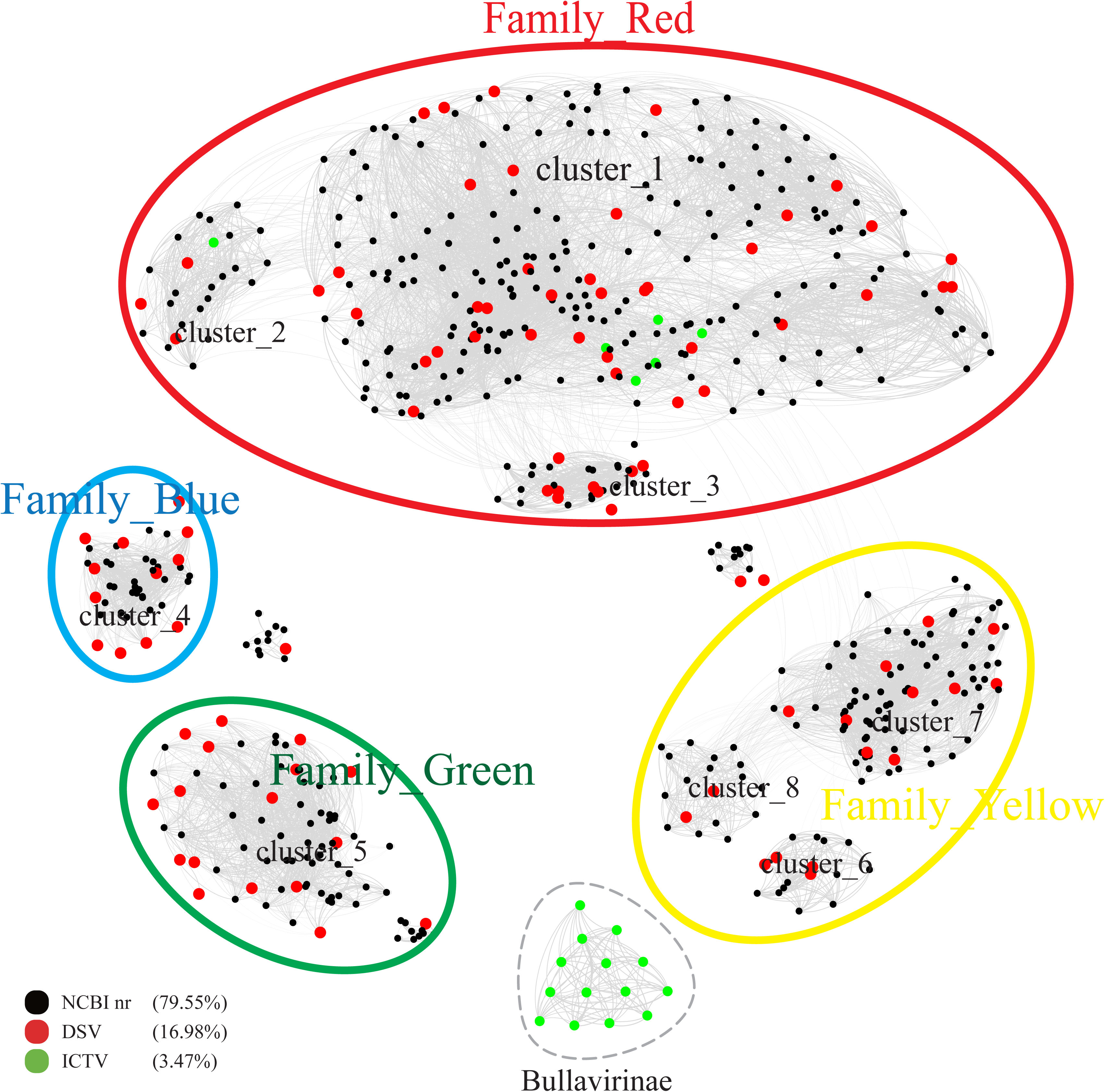
Similarity clustering network of Cap of microviruses from poultry slaughterhouses and related microvirus groups. The network includes identified microvirus Cap sequences from the DSV data (n=98), along with related Cap sequences from the NR data (n=459) and microvirus Cap sequences from the ICTV data (n=20). The similarity clustering network was constructed using Gephi (version 0.9.7) based on Diamond (version 0.9.14.115) alignment score. Gray connections represent Diamond Blastp score >480.

## 3 Results

### 3.1 Discovery of Novel Subfamilies of Microviridae

According to the ICTV standards, Microviridae includes two subfamilies (Bullavirinae and Gokushovirinae) and seven described genera(15). Among them, the subfamily Bullavirinae has three genera, comprising 14 species. The subfamily Gokushovirinae has four genera, consisting of eight species. We selected 98 complete DNA viral genomes from the DSV dataset, annotated as Microviridae, with a genome integrity exceeding 90%, for in-depth analysis. All DSV genomes have lengths ranging from 4 - 6 kb, consistent with the genome size of Microviridae(13). As predicted, these viruses all have Cap with lengths of 450-600 amino acids (AA). According to the Cap similarity clustering network graph (Figure 1), the Microviridae sequences from DSV, along with the related sequences aligned in NR and the Microviridae sequences from ICTV (totaling 577 sequences), roughly cluster into 9 clusters (cluster_1 to 8 and Bullavirinae). Among them, the 14 sequences of Bullavirinae (14 distinct species recognized by ICTV) formed a separate cluster (light green). It is worth noting that cluster_1 (C1) and cluster_2 (C2) include an additional 6 ICTV sequences, all of which belong to Gokushovirinae. In C1, there are 5 ICTV sequences, 4 of which belong to Chlamydiamicrovirus, and 1 belongs to Spiromicrovirus. In C2, 1 ICTV sequence belongs to Bdellomicrovirus. According to this classification standard, the other unclassified viruses in C1 and C2 should also belong to the Gokushovirinae. The remaining 6 clusters (cluster_3 to 8) do not include ICTV sequences, suggesting that these clusters might represent novel subfamilies within the Microviridae.

Based on the similarity of these 9 clusters, we can further categorize them into 5 major families, tentatively referred to as Family_Red, Family_Blue, Family_Green, Family_Yellow, and the independent Bullavirinae cluster. Among them, Family_Yellow includes 3 clusters, namely cluster_6 (C6), cluster_7 (C7), and cluster_8 (C8). Family_Red includes C1, C2, and cluster_3 (C3), which contain 6 Gokushovirinae ICTV sequences. Therefore, for now, we equate Family_Red with the Gokushovirinae. However, Family_Blue, Family_Green, and Family_Yellow do not include any ICTVsequences, suggesting that they are newly discovered taxonomic units in this study, at least on par with Gokushovirinae and Bullavirinae. Since Microviridae is already classified at the family level, whether Family_Blue, Family_Green, and Family_Yellow are proposed as new subfamilies or families requires further discussion.

### 3.2 Expanding the Potential Hosts of Microviruses

Host prediction was performed on the 98 newly discovered microvirus sequences from this study, along with their associated 459 NR sequences and 20 ICTV sequences, using hostG(37) and cherry(38). In the hostG results, only NC_002643.1 from ICTV was accurately predicted to have a host (Bdellovibrio). However, in the cherry results, the majority of hosts were consistent with the ICTV results, indicating that the success rate and accuracy of cherry predictions were higher than hostG. According to the hostG results, the main hosts for DSV were Bdellovibrio and Chlamydia (Figure 2a, b). For NR sequences, the hosts were mainly distributed in the Bdellovibrio, Chlamydia, and Parabacteroides. Although these results align well with the current understanding of microvirus hosts, results from cherry suggest (Figure 2c, d) that the hosts of microviruses may be far more diverse than these three genera.

**Figure 2.**
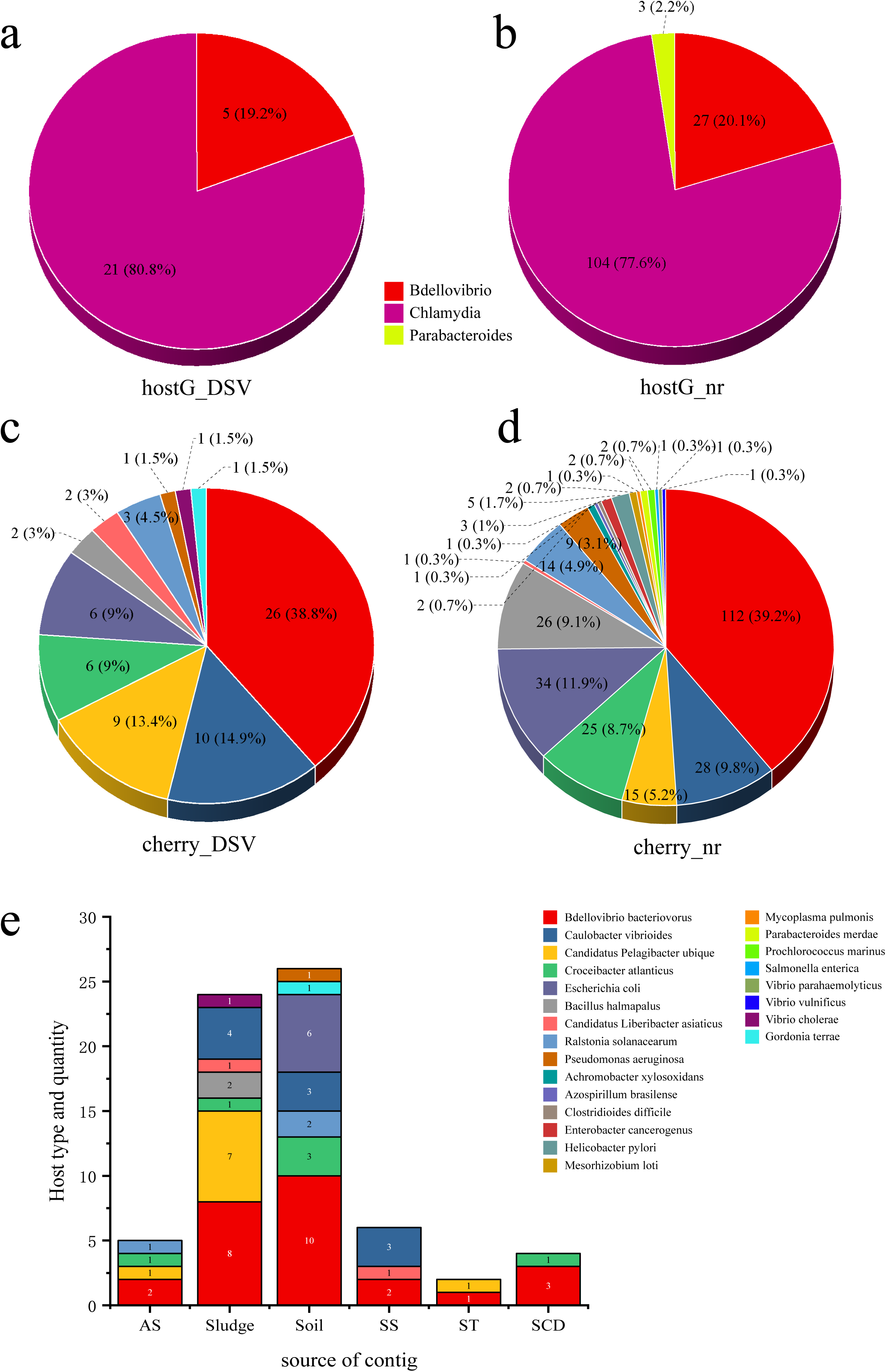
The host types and quantity statistics of microviruses from poultry slaughterhouses and related groups. (a) hostG(37) results of DSV sequences. (b) hostG results of NR sequences. (c) cherry(38) results of DSV sequences. (d) cherry results of NR sequences. Score > 0.7. (e) Host types and quantity predicted by cherry for corresponding DSV sequences. AS (All soil and sludge of slaughterhouse); Soil (Soil of slaughterhouse); Sludge (Sludge of slaughterhouse); SS (Swab of slaughterhouse workshop); ST (Swab of poultry transport vehicle); SCD (Oral and cloacal swabs of chickens and ducks).

Although Bdellovibrio bacteriovorus and Escherichia coli are the predominant hosts in the NR-derived virus hosts, respectively. Cherry also predicted hosts such as Caulobacter vibrioides and Bacillus halmapalus, indicating numerous microvirus hosts that have not been previously reported. Among the DSV-derived virus hosts, Bdellovibrio bacteriovorus still dominated, followed by Caulobacter vibrioides and Candidatus Pelagibacter ubique, representing novel hosts. In addition, NR data revealed the presence of human and animal pathogens such as Helicobacter pylori and Enterobacter cancerogenus. The DSV data also identified potential hosts including Vibrio cholerae and Pseudomonas aeruginosa. This suggests that the host range of microviruses within the Microviridae may be extensive, and that there are likely more potential hosts yet to be discovered.

From the perspective of sample types, the highest abundance of microviruses was observed in soil and sludge samples, corresponding to a higher diversity and quantity of their respective hosts (Figure 2e). Bdellovibrio bacteriovorus, as a typical host for microviruses, showed a higher proportion across various samples. Caulobacter vibrioides also exhibited high abundance in sludge, soil, and the slaughterhouse workshop (Figure 2e). While Escherichia coli, Gordonia terrae, and Pseudomonas aeruginosa were predicted only in soil samples, Vibrio cholerae was exclusively found in sludge samples. Other host bacteria were detected across different sample types. This indicates a close relationship between the detection of microviruses and the distribution of their host bacteria, displaying certain characteristics in various sample types.

### 3.3 Genome Length and GC Content

The genome sizes and GC content of viruses within the same family or genus are usually relatively consistent(45, 46). Based on the identification of 9 clusters in the previous sections, we further created boxplots illustrating their genome size and GC content (Figure 3). Both genome size and GC% exhibited high consistency within each of the 9 clusters, while significant differences were observed among different clusters. For instance, Bullavirinae showed distinct genome sizes and GC content compared to other groups. Individual scattered black dots outside the boxes in the figure represent sequences from the NR data. These results indicate that the genome characteristics of microviruses from different taxonomic groups exhibit good consistency and indirectly validate the reliability of our classification method based on the similarity clustering network graph.

**Figure 3.**
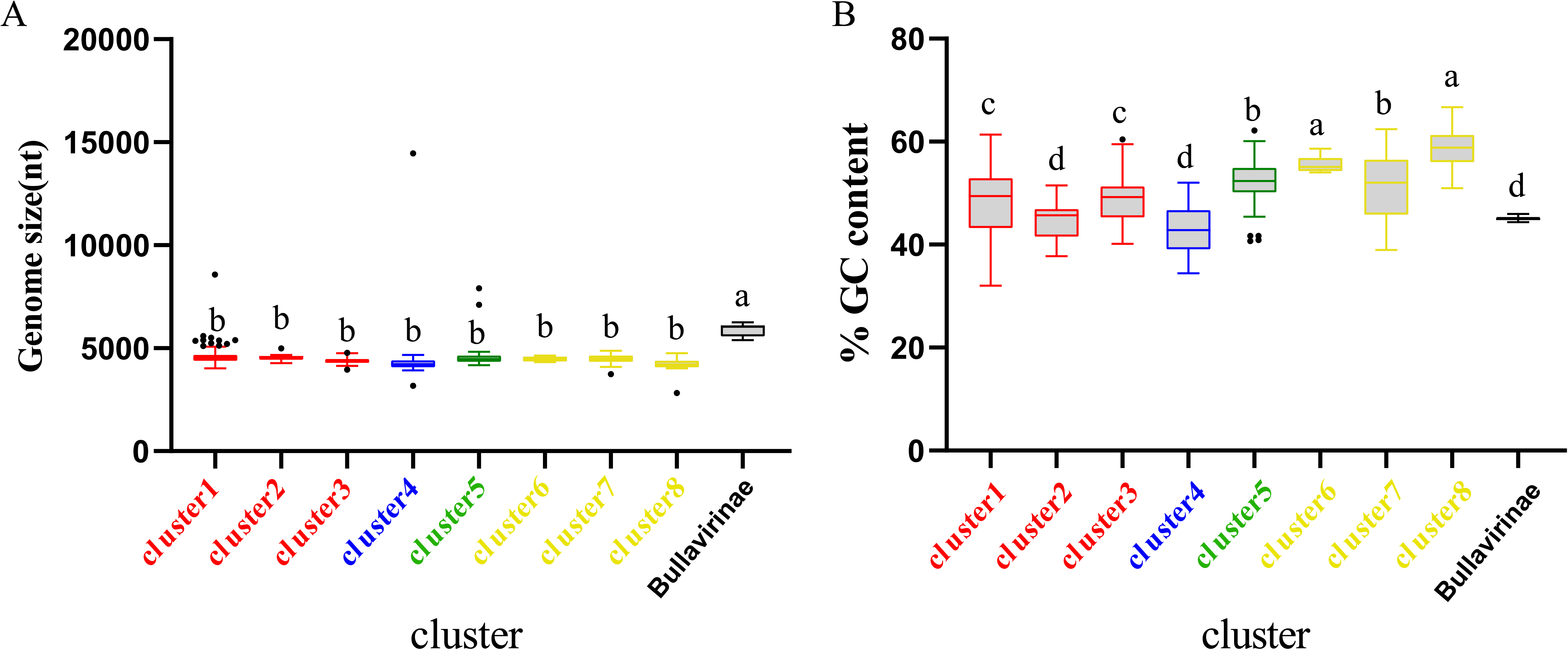
Genomic features of each cluster of microviruses from poultry slaughterhouses and related groups. (A) Distribution of microviruses genome sizes in each cluster from Result 3.1. (B) Distribution of microviruses genome %GC content in each cluster from Result 3.1. Red, blue, green, yellow, and black correspond to Family_Red, Family_Blue, Family_Green, Family_Yellow, and Bullavirinae, respectively. Turkey’s test was used, where P < 0.05 indicates significant differences, and P > 0.05 indicates no significant differences. In the group where the maximum mean value is located, mark it with the letter “a.” Then, compare this mean value with the mean values of other groups one by one. If there is no significant difference, label them with the same letter “a.” Continue this process until encountering a mean value with a significant difference, then label it with the letter “b.” Subsequently, use “b” as the standard for further comparisons. Repeat this process, labeling consecutive mean values with the letters “b” until encountering a mean value with a significant difference, which is then labeled as the letter “c.” This pattern continues for subsequent comparisons. The plot displays median values, 25th and 75th percentiles, 1.5 interquartile ranges, and outlier data points.

### 3.4 Phylogenetic Analysis Based on Cap Sequences

To better illustrate the diversity of DSV-related microviruses and their evolutionary origins, phylogenetic trees were constructed for each cluster based on the results in Figure 1. The sequences of DSV-related microviruses were classified and named according to the sample source and host type of the viruses (see Materials and Methods 2.10 for reference).

C1 is the cluster with the highest number of members among the eight clusters and exhibits the most diverse range of host sources (Figure 4. Displayed are partial positions of the phylogenetic tree. Some phylogenetic branches have been collapsed, the complete phylogenetic tree is detailed in Figure S1). Notably, in addition to typical hosts such as Escherichia coli and Bdellovibrio bacteriovorus, this cluster has hosts that were previously unreported, such as Caulobacter vibrioides, Pseudomonas aeruginosa, and Helicobacter pylori. Caulobacter vibrioides is a Gram-negative oligotrophic bacterium widely distributed in freshwater lakes and streams, serving as an important model organism for studying cell cycle regulation, asymmetric cell division, and cell differentiation(47). Pseudomonas aeruginosa is a common multidrug-resistant pathogen, characterized by its capsule, Gram-negative nature, and aerobic or facultatively anaerobic growth, causing diseases in plants and animals, including humans(48). Helicobacter pylori is a Gram-negative, flagellated spiral bacterium, classified as a class I carcinogen, responsible for approximately 89% of gastric cancer cases and associated with 5.5% of cancer cases worldwide(49–51). In general, hosts within the same clustering branch are relatively homogeneous. For example, in Figure 4, the hosts in the purple-colored block branch are primarily Bdellovibrio bacteriovorus, while the hosts in the deep blue-colored block branch are mainly Caulobacter vibrioides.

**Figure 4.**
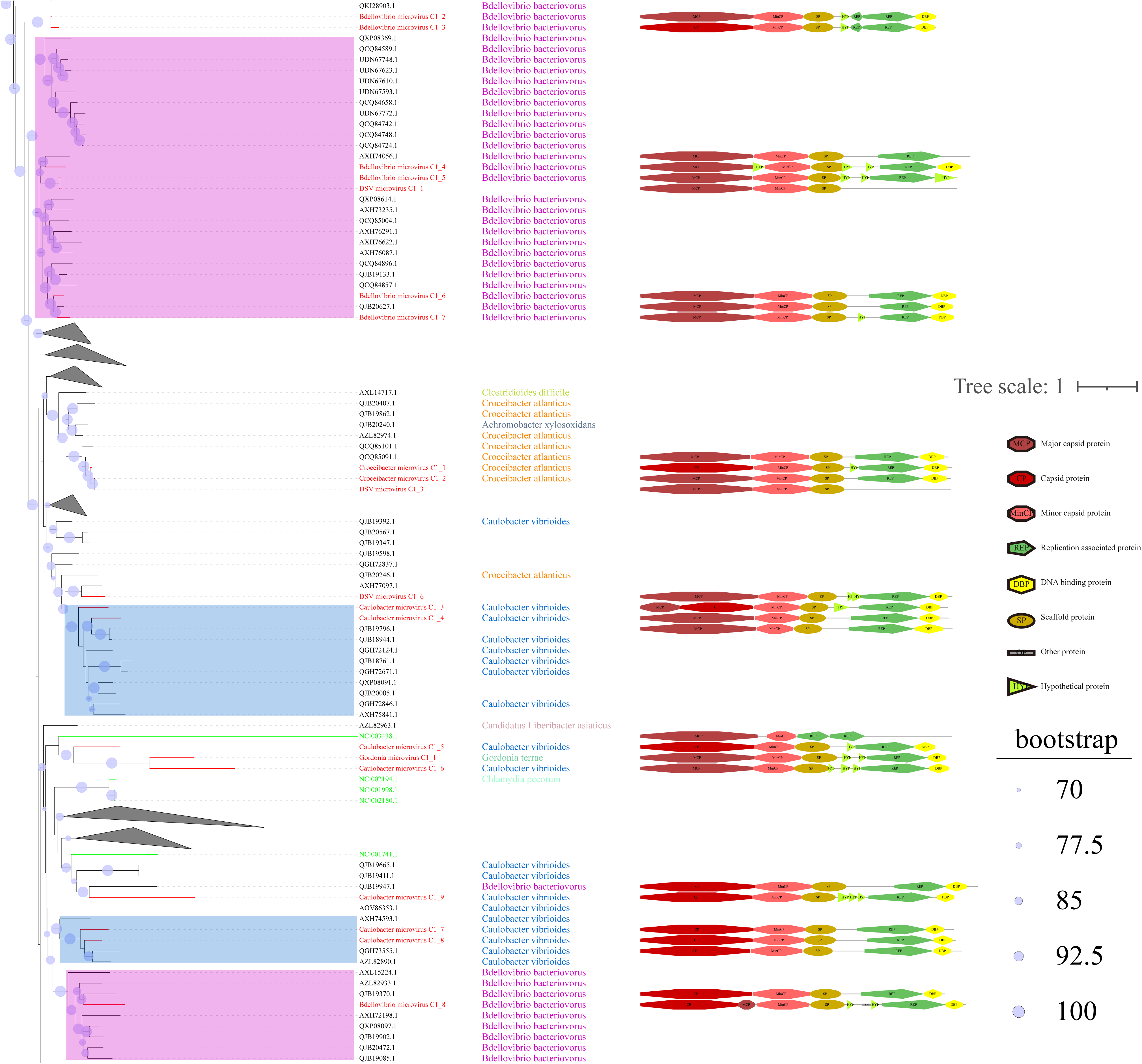
Phylogenetic tree, hosts, and genomic structure of cluster_1 microviruses from poultry slaughterhouse and related sources. The maximum likelihood phylogenetic tree was constructed based on the Cap sequences of Microviruses using IQtree (version 2.1.4). ModelFinder was set to MFP, and 1000 ultrafast bootstrap replicates were performed, displaying bootstrap values > 70. The red branches represent DSV microvirus sequences, green branches represent ICTV sequences, and black branches represent NR sequences. The third column shows host annotations predicted by Cherry, and the fourth column displays partial genomic structure diagrams.

From the sample sources perspective, DSV viruses in this cluster mainly originate from soil (CD-SXG) and sludge (CD-SXD), with a few from swab samples taken in the slaughterhouse workshop environment (CD-SXS) (Supplementary Table S1). In comparison, the sources of NR viruses are more diverse, including animal metagenomes, wastewater metagenomes, human metagenomes, and blackflies. As seen in the genomic structure diagram in Figure 4, members of C1 typically possess signature genes such as Major capsid protein or capsid protein(Cap), and Replication associated protein (Rep). Moreover, the genomes in this cluster often exhibit a sequential arrangement of Cap, Minor capsid protein (MinCP), Scaffold protein(SP), Rep, and DNA binding protein (DBP). However, a few viruses in this cluster have genome organization sequences that deviate from this pattern, such as DSV microvirus C1_4. Furthermore, Replication-associated protein was not predicted in DSV microvirus C1_1 and DSV microvirus C1_3. Coincidentally, these two sequences also lack host prediction results, likely suggesting the novelty of these viral genomes. Overall, sequences with closer phylogenetic relationships tend to exhibit more apparent consistency in host specificity and genomic structure.

C2 is a small viral cluster with a consistent host source, all being Bdellovibrio bacteriovorus, and a highly consistent genomic structure (Figure 5). The viruses in this cluster exhibit a sequential arrangement of Cap, MinCP, SP, DBP, and Rep, with Hypothetical protein (HYP) inserted on either side of DBP. Bdellovibrio microvirus C2_2 and Bdellovibrio microvirus C2_1 are closely related to AZL82867.1 and QJB19506.1, respectively. The genome of AZL82867.1 is derived from Honey bees, while QJB19506.1 originates from wastewater metagenome. This observation suggests the widespread presence of microviruses and their Bdellovibrio bacteriovorus hosts in various environmental settings. Only three sequences in C3 were predicted to have hosts (Figure S2), indicating that this group lacks sufficient host information and is a relatively novel group compared to C1 and C2. The predicted hosts for these three sequences are Azospirillum brasilense (A. brasilense) and Enterobacter cancerogenus (E. cancerogenus). A. brasilense is a microaerophilic nitrogen-fixing bacterium widely present in the rhizosphere worldwide, promoting plant growth(52, 53). E. cancerogenus is a significant pathogen commonly found in human clinical specimens such as blood and cerebrospinal fluid. It is not sensitive to penicillin and cephalosporin(54), and exploring bacteriophage targeting such multidrug-resistant pathogens is meaningful for developing phage therapy methods.

**Figure 5.**
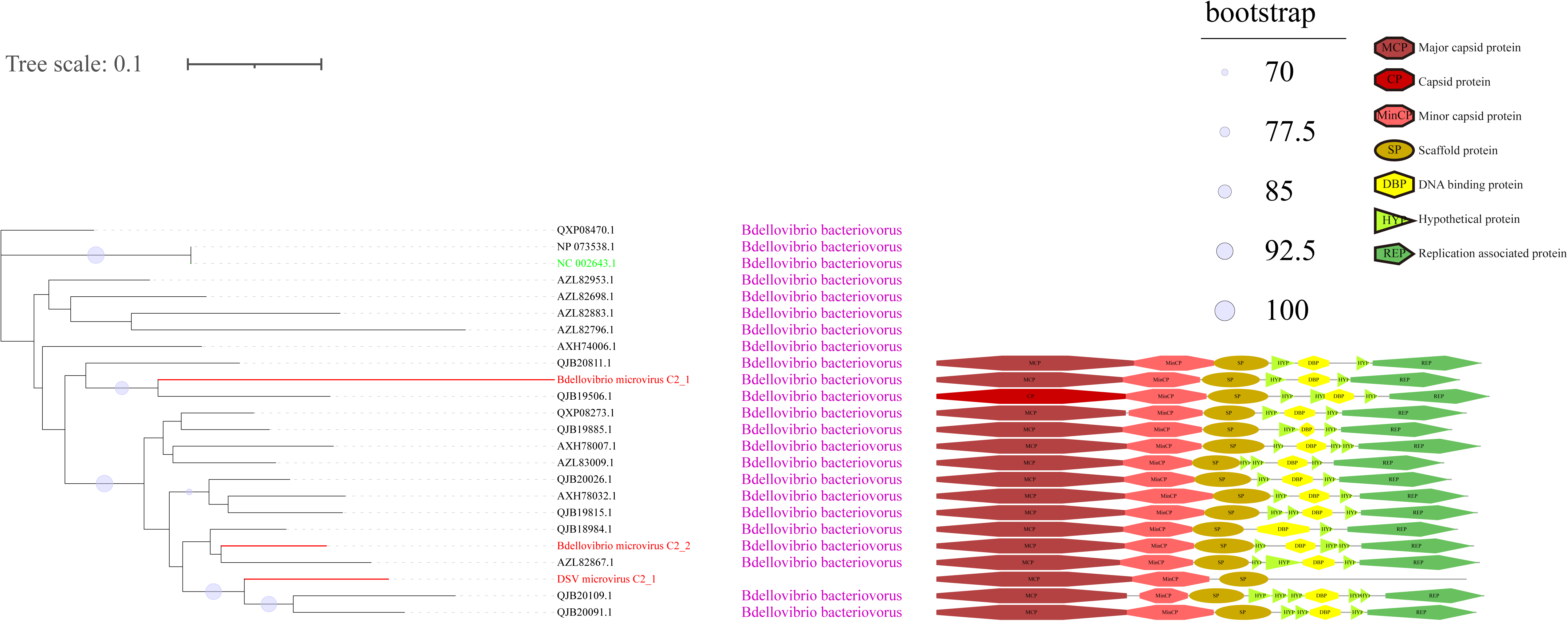
Phylogenetic tree, hosts, and genomic structure of cluster_2 microviruses from poultry slaughterhouse and related sources. The maximum likelihood phylogenetic tree was constructed based on the Cap sequences of Microviruses using IQtree (version 2.1.4). ModelFinder was set to MFP, and 1000 ultrafast bootstrap replicates were performed, displaying bootstrap values > 70. The red branches represent DSV microvirus sequences, green branches represent ICTV sequences, and black branches represent NR sequences. The third column shows host annotations predicted by Cherry, and the fourth column displays partial genomic structure diagrams.

Cluster_4 (C4) generally exhibits a relatively tidy genome structure (Figure S3). It is noteworthy that, despite being in the same cluster, there are significant differences in the host sources between DSV and NR viruses in C4. The majority of NR viruses are derived from wastewater metagenome samples, and their hosts are predominantly Bacillus halmapalus. Bacillus halmapalus, a halophilic bacterium, is a Gram-positive, alkaliphilic, alkalitolerant, facultative anaerobe. It is commonly isolated from soil, and its pathogenicity is not well understood(55). In DSV, only two viruses have Bacillus halmapalus as their host, and both are derived from sludge samples, aligning with the source of this bacterial species. Unlike NR, the primary hosts for DSV viruses are Candidatus Pelagibacter ubique. Except for Candidatus Pelagibacter microvirus C4_1, which originates from a swab of the transportation vehicle (CD-SXt), the rest are all from sludge samples (CD-SXD) (Table S1). Studies suggest that Candidatus Pelagibacter species may be among the most abundant bacteria globally and play a crucial role in the carbon cycle on Earth.

Croceibacter atlanticus belongs to the phylum Bacteroidetes and is a species isolated from the Atlantic Ocean(56). Croceibacter microvirus C4_1 is the only virus in this cluster derived from swab of duck oral and cloaca (D-SXAO) (Table S1), and it is specifically associated with the host Croceibacter atlanticus. This observation once again confirms the conclusion from Figure 2e that the detection of microviruses is closely related to the distribution of their host bacteria and the source of the samples.

Cluster_5 (C5), as shown in Figure S4. Although most NR sequences include Cap and Rep, DSV sequences, such as Bdellovibrio microvirus C5_2/4/7, Escherichia microvirus C5_1/2/13, only predict 2 ORFs: capsid protein and MinCP. We have not observed a correlation between this situation and sample sources, indicating a potentially higher novelty and lower conservation of genes in DSV sequences. C6 (Figure S5), similar to C3 (Figure S2), is a smaller cluster without predicted hosts, indicating a need for further research on this cluster. The genomic structure of the C7 sequences is primarily arranged in the order of Cap, MinCP, SP, Rep, and DBP (Figure S6). Croceibacter microvirus C7_1 and Croceibacter microvirus C7_2, two viruses within the same major branch, share Croceibacter atlanticus as their host (previously introduced in C4). This branch is the only one with predicted host results, while other branches lack host predictions. Therefore, C7 is also a potentially interesting virus cluster worthy of in-depth research. C8 (Figure S7) has two distinct hosts, with Ralstonia solanacearum being the dominant host and Achromobacter xylosoxidans as the second host. Ralstonia solanacearum is considered one of the most important plant pathogens due to its lethal nature, persistence, wide host range, and extensive geographical distribution(57). Achromobacter xylosoxidans belongs to the genus Achromobacter and is commonly found in moist environments, causing diseases such as bacteremia, pneumonia, pharyngitis, and urinary tract infections(58, 59).

### 3.5 Comparing DSV Viruses in Microvirus’s Virosphere

To better understand the relationship between the identified microviruses in the poultry slaughterhouse and other reported microviruses in the Microviridae family, we expanded our focus beyond the 577 viral genomes highlighted in this study (Figure 1). To this end, an additional set of 4077 microvirus Cap sequences (utilizing 4,007 sequences for this study) studied by Paul et al.(36) were incorporated into our analysis for a more comprehensive clustering analysis. In the study by Paul et al., microviruses were broadly classified into 19 families, corresponding to the 19 color-coded clusters in Figure 6. Upon comparing the clustering results between Figure 1 and Figure 6, there is a good overall agreement between the two figures. Specifically, the four families identified in Figure 1 are concentrated within the purple cluster in Figure 6, representing the largest cluster in Family 3, as defined by Paul et al. DSV viral sequences are predominantly distributed within Gokushovirinae A(36), Shukshmavirinae(60) and Group D(61) of Family 3. Additionally, three scattered sequences are found in Pichovirinae(14), Gokushovirinae B and Gokushovirinae C(36), indicating that microviruses in the poultry slaughterhouse environment primarily belong to these groups. This result suggests that, although microviruses in the poultry slaughterhouse environment exhibit high diversity and novelty, they may still be relatively underrepresented in the entire microvirus virosphere. Family 3, possibly due to its close association with the human environment, is the largest group within the Microviridae. The expansion of other groups awaits further supplementation with samples from different sources and microbial hosts.

**Figure 6.**
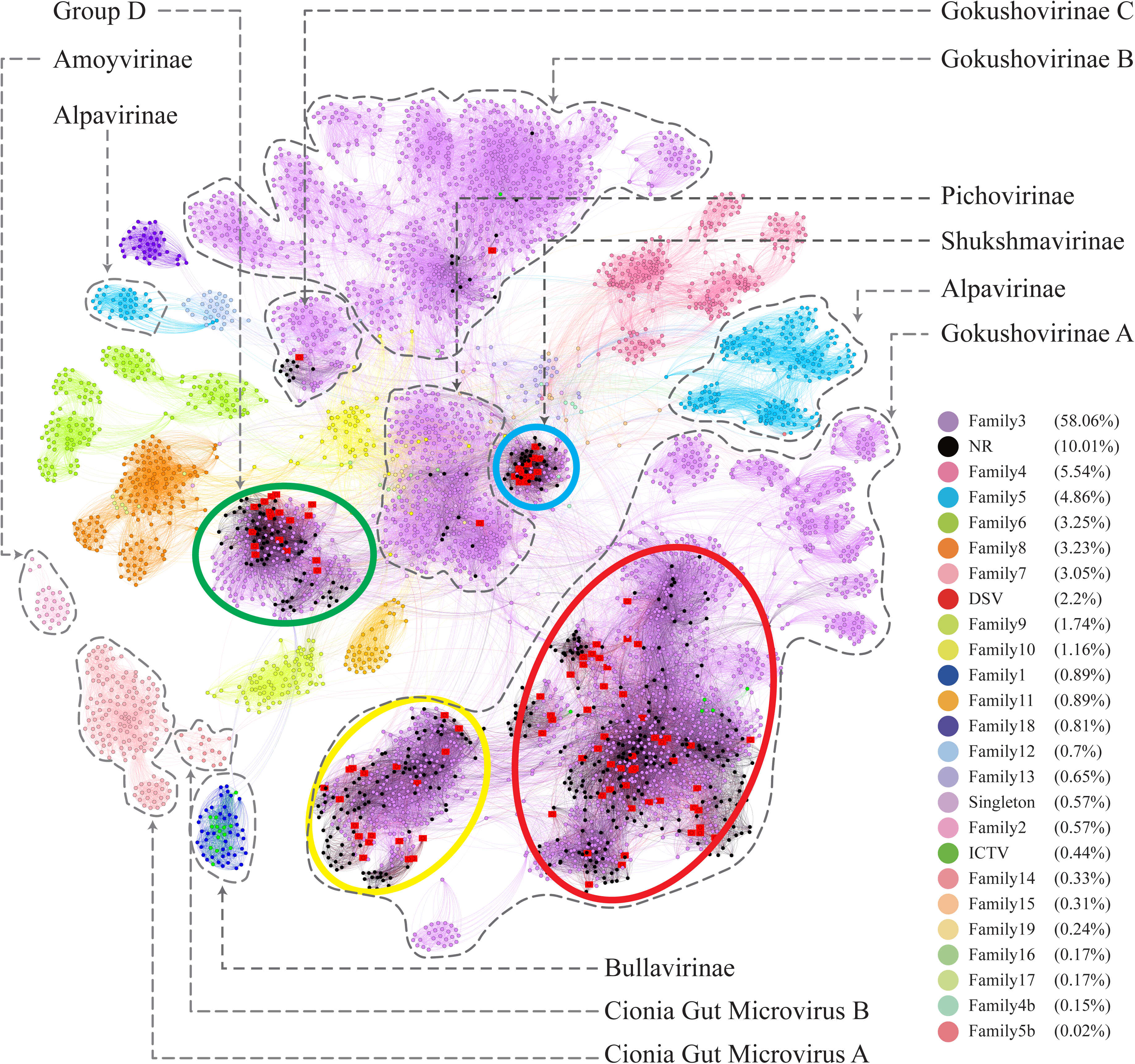
The microviruses collection with diverse taxa. Similarity clustering network was constructed using microviruses Cap sequences identified from DSV data (n=98), along with related Cap sequences from NR (n=459), ICTV microviruses Cap sequences (n=20), and an additional set of Cap sequences reported by Paul et al. (n=4007) (red dots represent DSV sequences, black dots represent NR sequences, and dark green dots represent ICTV sequences; other dots are colored based on the families defined by Paul et al.(36)). Clusters corresponding to those in Figure 1 are enclosed by ellipses of four different colors. Labels such as Pichovirinae, Shukshmavirinae, Group D, Alpavirinae, Gokushovirinae A/B/C correspond to subfamilies reported by previous studies(14, 16, 60, 61) and suggested classifications by Paul et al.(36) The similarity clustering network graph was created using Gephi (version 0.9.7) based on Diamond (version 0.9.14.115) alignment score, with gray edges indicating Diamond Blastp score >0.

On the other hand, according to the clustering results in Figure 6, the 19 major families delineated by Paul et al. can be further subdivided into approximately 45 smaller clusters. Particularly, within family 3 (the purple clusters), our clustering method can split it into as many as 20 smaller clusters. Specifically, Group D, Shukshmavirinae, Pichovirinae, and Gokushovirinae C each form independent cluster, while Gokushovirinae A and Gokushovirinae B can be further divided into multiple clusters at the same hierarchical level. These clusters are at least equivalent in status to Group D, Shukshmavirinae, and Pichovirinae. Additionally, families of other colors can also be further subdivided into smaller clusters. For example, family 5 identified as Alpavirinae(16) can be distinctly clustered into 4 different clusters in this study. This suggests that these smaller clusters may represent novel subfamilies or families that require further identification.

### 3.6 The Relationship between the Clusters of Microviruses and Their Host Sources

According to the cherry(38) results, the points in Figure 1 and Figure 6 are colored coded on host types in Figure 7. As shown in Figure 7a, clusters C2, C4, C5, a portion of C7, C8, and the Bullavirinae cluster exhibit clear host specificity, while the host colors in cluster C1 are highly mixed. Specifically, Bdellovibrio bacteriovorus is mainly the host for C2 and C5, the hosts for C4 are primarily Bacillus halmapalus and Candidatus Pelagibacter ubique, the main host for C8 is Ralstonia solanacearum, and only some sequences in C7 have host results, all of which are associated with Croceibacter atlanticus. Cluster C1 includes a significant number of Bdellovibrio bacteriovorus viruses, as well as viruses infecting Escherichia coli, Caulobacter vibrioides, Croceibacter atlanticus, and other bacteria. This may suggest that this group of viruses is more prone to host jumping compared to other viruses group.

**Figure 7.**
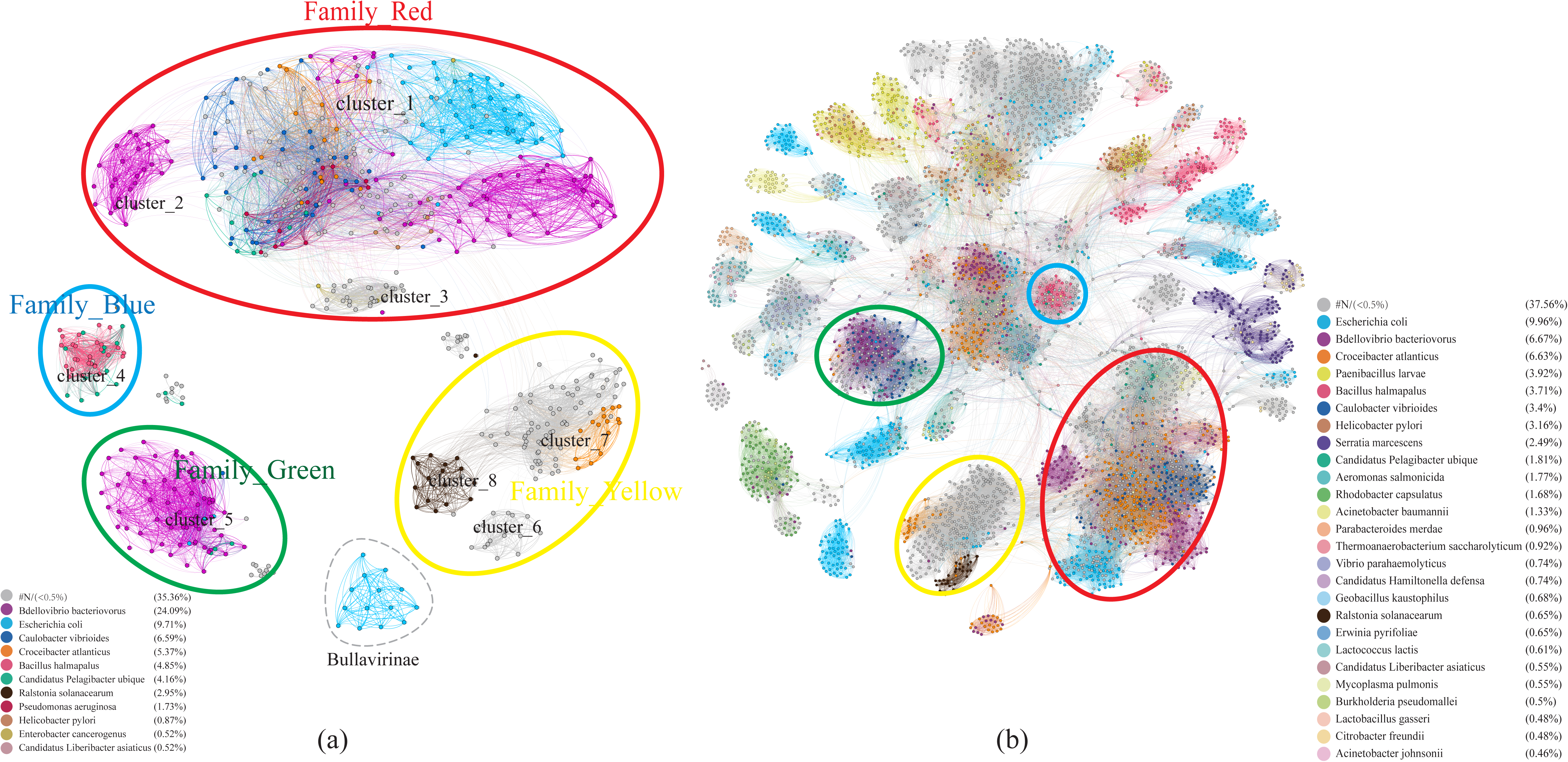
Host specificity of different clusters of Microviruses. (a) Similarity clustering network constructed using microviruses Cap sequences identified in DSV data (n=98), along with related Cap sequences from NR (n=459), and microviruses Cap sequences from ICTV (n=20), colored based on host types. (b) Based on the sequences in (a), an extended similarity clustering network was constructed by introducing Cap sequences reported by Paul et al. (n=4007), also colored according to host types. The similarity clustering network graph was created using Gephi (version 0.9.7) based on Diamond (version 0.9.14.115) alignment score, with gray edges indicating Diamond Blastp score >0.

Similar to the results in Figure 7a, the host sources of Family_Red members in Figure 7b remain diverse, primarily involving Bdellovibrio bacteriovorus, Caulobacter vibrioides, Croceibacter atlanticus, Escherichia coli. As the number of members increases, Family_Green shows an expanded range of host types, mainly associated with Caulobacter vibrioides, Bdellovibrio bacteriovorus, and Candidatus Pelagibacter ubique. Family_Blue continues to be dominated by Bacillus halmapalus and Candidatus Hamiltonella defensa. Although the number of Family_Yellow members has increased significantly, a majority still lacks predicted hosts. Apart from the four main groups focused on in this study, the host types of most smaller groups are relatively singular, such as Bacillus halmapalus (family4), Escherichia coli (family5), Rhodobacter capsulatus (family7) (Figure 7b). These results suggest variations in host specificity among different viral groups. Moreover, the good correspondence between similarity clustering networks and host prediction results is evident.

## 4 Discussion

Viruses are the most abundant life forms on earth. It is estimated that there are as many as 10^31^ virus particles on earth(11, 12). However, The International Committee on Taxonomy of Viruses (ICTV) has officially recognized only around 12,000 known virus species (https://ictv.global/vmr, Version: VMR MSL38 v1). Viruses are often considered the “dark matter” of life sciences. Due to the challenges in cultivating many viruses, our understanding is limited to those that are easily cultivated and have significant impacts on humans or the economy. Advances in high-throughput sequencing and virome technologies have overcome the dependency on host cell cultures in traditional virology research, greatly enhancing the efficiency of discovering and identifying new viruses(62). In recent years, virome technologies has been widely applied in various studies, including marine environments and research on vertebrates and invertebrates, leading to the identification of numerous novel viruses(63, 64) and significantly expanding our knowledge of the viral world.

Microviridae is one of the most common families of single-stranded DNA (ssDNA) viruses. Compared to double-stranded DNA (dsDNA) phages, the genomes of Microviridae are smaller, typically exhibiting higher safety by being less prone to carry virulence and resistance genes(36). Moreover, they are widely distributed across various ecosystems(65, 66), representing a relatively accessible and exploitable source of DNA resources. As of now, the ICTV recognizes only two subfamilies within Microviridae, namely Bullavirinae and Gokushovirinae. However, this classification does not fully capture the extensive diversity of newly reported microviruses(36). In recent years, numerous new taxonomic groups within the Microviridae family have been proposed. For instance, Krupovic et al. introduced a novel Microviridae subfamily named Alpavirinae, which was identified as prophage(16). Additionally, several newly proposed subfamilies of Microviridae include Pichovirinae from the human gut(14), Sukshmavirinae from termites(60), Group D from dragonflies(61), and Aravirinae and Stokavirinae from sphagnum-dominated peatlands(39). Paul et al. comprehensively analyzed the genomes of microviruses using their classification method, providing insights into the diversity, distribution, and host range of this viral group(36). The proposed new classifications are clearly represented in the clustering network graph of this study (Figure 6), indicating a good validation across different research efforts.

As far as we know, this study represents a relatively comprehensive compilation of members of the Microviridae, providing an overview of the classification of Microviridae and holding significant importance for the identification, exploration, and expansion of Microviridae. However, due to the large number of potential new taxa, this study did not assign explicit taxonomic names to them, focusing instead on demonstrating relationships between clusters. We believe that as more members of Microviridae are discovered and identified, this family will continue to give rise to new taxa and may undergo redefinition. To address this situation, there is an urgent need for a universal and straightforward method for classification, such as utilizing numbers or letters to systematically name newly emerging taxa.

The evolutionary trajectory of dsDNA phages is primarily influenced by horizontal gene exchange, driving the diversity and adaptive evolution of this phage class. However, for ssDNA phages, the evolutionary patterns may fundamentally differ(67–69). For instance, in microviruses, gene recombination is not widespread, and the presence of Cap may limit the insertion of foreign DNA sequences(70), potentially restricting gene transfer at the horizontal level. Despite these factors, microviruses exhibit high mutation rates in their genomes(71), suggesting that they may employ different evolutionary strategies to enhance adaptability. This adaptability is evident in the diverse clusters and extensive host range discovered in this study, as well as the diversity in hosts and genome structures found even within the same phylogenetic branch. Additionally, both this study and others(14) have observed differences in the genome structures of microviruses from various sample sources or types. This reflects the complexity and diversity of their evolution, showcasing their ability to adapt to different environments.

At present, the mainstream view suggests that the hosts of microviruses are primarily intracellular parasitic bacteria and Enterobacteria. For instance, the host of the Bullavirinae is Enterobacteria, and detailed studies have been conducted on representatives of this family, such as the phage ΦX174(69). Members of the the Gokushovirinae only infect Chlamydia, Bdellovibrio and Spiroplasma(13, 72). However, an increasing number of studies indicate that microviruses can infect a broader range of bacterial hosts, including Vibrio parahaemolyticus(73, 74), Salmonella(75), Shigella flexneri(76) and other bacteria. To address the question of infecting hosts, this study employed two new host prediction methods. The hostG utilizes shared protein clusters between viruses and prokaryotes to create a knowledge graph and trains a graph convolutional network for prediction(37). While it achieves high prediction accuracy, its results tend to be conservative and can only predict hosts at the genus level. Cherry is described as having the highest accuracy in identifying virus-prokaryote interactions, outperforming all existing methods at the species level, with an accuracy of 80%(38). The results from Cherry indicate that the hosts of the Microviridae exhibit extremely high diversity, including various pathogens such as Mycoplasma pulmonis, Helicobacter pylori, Vibrio cholerae, Clostridioides difficile, and Pseudomonas aeruginosa. Additionally, this study identified some plant-pathogenic bacteria, such as Ralstonia solanacearum (R. solanacearum) and Candidatus Liberibacter asiaticus (CLas). Bacterial wilt, caused by R. solanacearum, is economically significant as it can infect over 250 plant species, including potatoes, tomatoes, and tobacco, causing substantial yield losses in tropical and subtropical regions(77, 78). CLas is the pathogen responsible for citrus Huanglongbing (HLB, also known as citrus greening disease)(79), a highly destructive disease threatening global citrus production. There has been limited research on Microviridae infecting plant-pathogenic bacteria, and the findings of this study suggest that Microviridae also holds potential for applications in the control of bacterial diseases in plants.

## 5 Conclusion

This study employed virome techniques to thoroughly explore potential members of Microviridae in a poultry slaughterhouse, successfully identifying and analyzing 98 novel and complete microvirus genomes. Based on the similarity of Cap proteins, it was discovered that these genomes represent at least six new subfamilies within Microviridae, distinct from Bullavirinae and Gokushovirinae, as well as three higher-level classification units. These new taxa exhibit obvious regularities in genome size, GC content, and genome structure, further highlighting the rationality of the classification method used in this study. Additionally, based on the 19 families classified by previous researchers for all microviruses, our approach divides microviruses into about 45 more detailed clusters, which may serve as a new standard for classifying Microviridae members. The current information on microviruses’ hosts remains limited, and this study significantly expands their host range. In addition to typical hosts such as intracellular parasitic bacteria and Enterobacteria, we identified over 20 potential new hosts, including important pathogens like Helicobacter pylori and Vibrio cholerae. Moreover, we revealed distinct host specific differences among different taxa. These new findings will contribute to a deeper understanding of the interactions between Microviridae and their hosts.

## 7 Author Contributions

**XKM**: Methodology, Validation, Formal analysis, Data Curation, Writing - Original Draft, and Visualization; **LBF:** Conceptualization, Methodology, Sample collection, Project administration, and Funding acquisition; **ZP, SXY, LC and LGF**: Data Curation, and Investigation; **CXD**: Writing - Review & Editing; **PJQ, QSP, YXQ and LMS**: Sample collection; **JJZ**: Conceptualization, Methodology, Writing - Original Draft, Writing - Review & Editing, Supervision, Project administration, and Funding acquisition; **YLH**: Conceptualization, Methodology, Resources, Writing - Review & Editing, Supervision, Project administration, and Funding acquisition.

All authors read and approved the final manuscript.

## Acknowledgments

This project was supported by the Natural Science Foundation of China (nos. 31872499 and 31972847) to Yuan LH and Jiang JZ; the Innovation Team Project of Guangdong Universities (No. 2022KCXTD017) to Yuan LH; the Central Public-Interest Scientific Institution Basal Research Fund, CAFS (nos. 2023TD44 and 2021SD05) to Jiang JZ. The funders had no role in the study design, data collection, and analysis, decision to publish, or manuscript preparation.

## **8** Data availability

The data set supporting the results of this article has been deposited in the National Center for Biotechnology Information (NCBI) under BioProject accession code PRJNA1053868. All viral genomes obtained in this study were deposited in GenBank with the accession numbers: OR998966-9063.

---

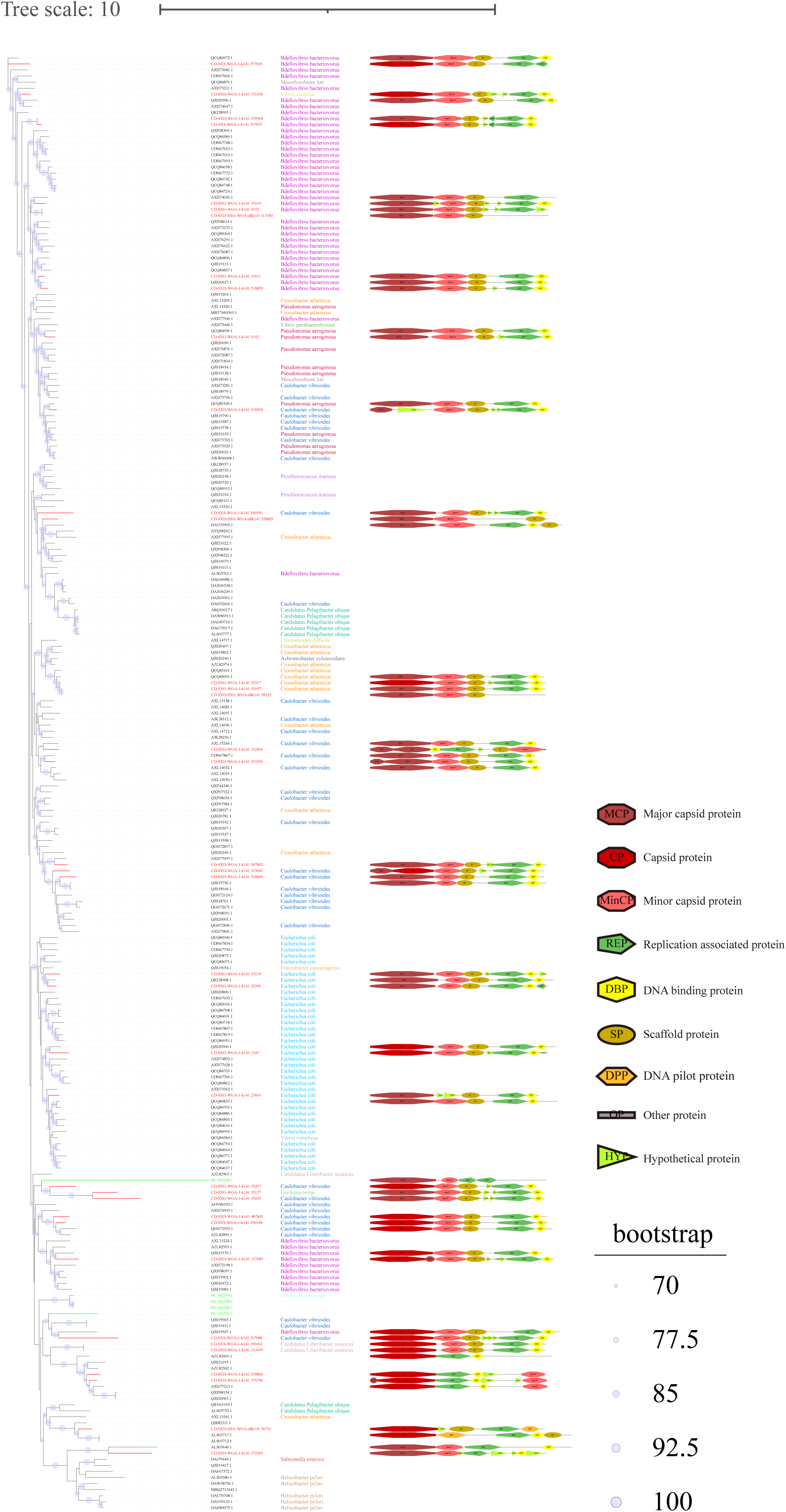

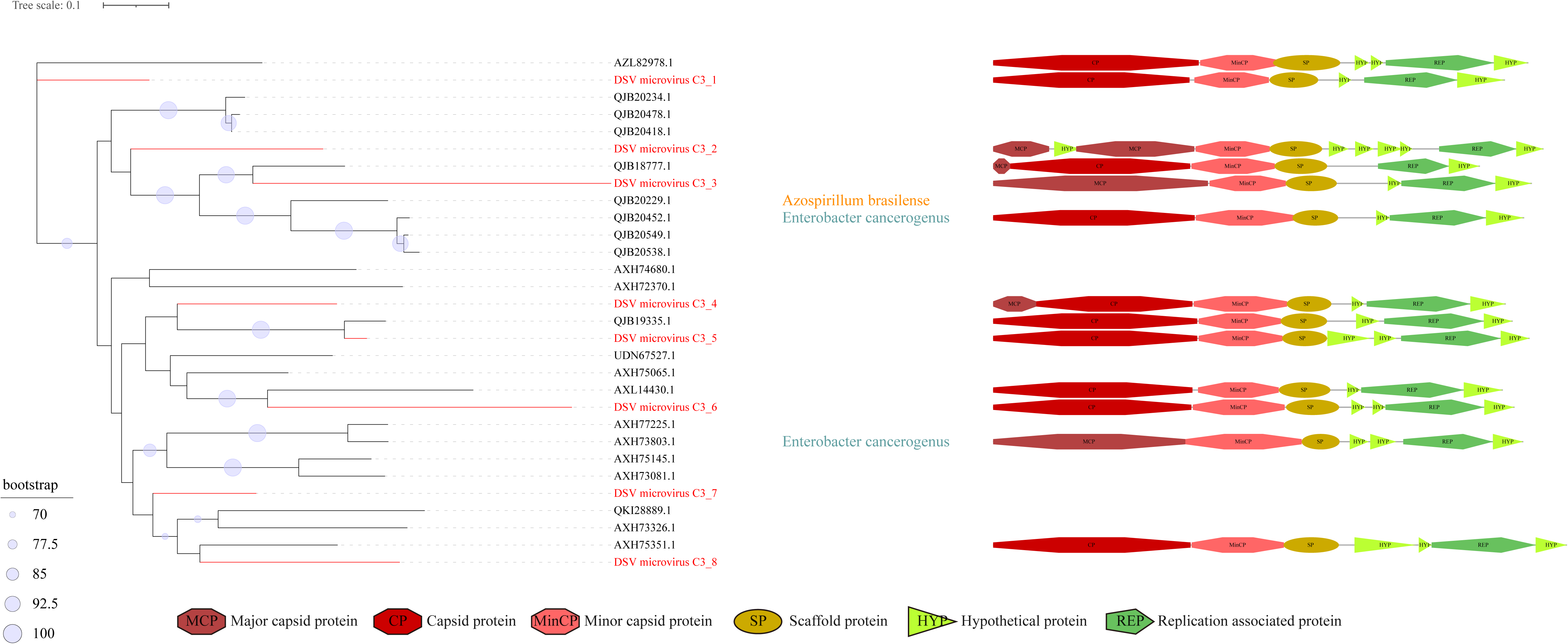

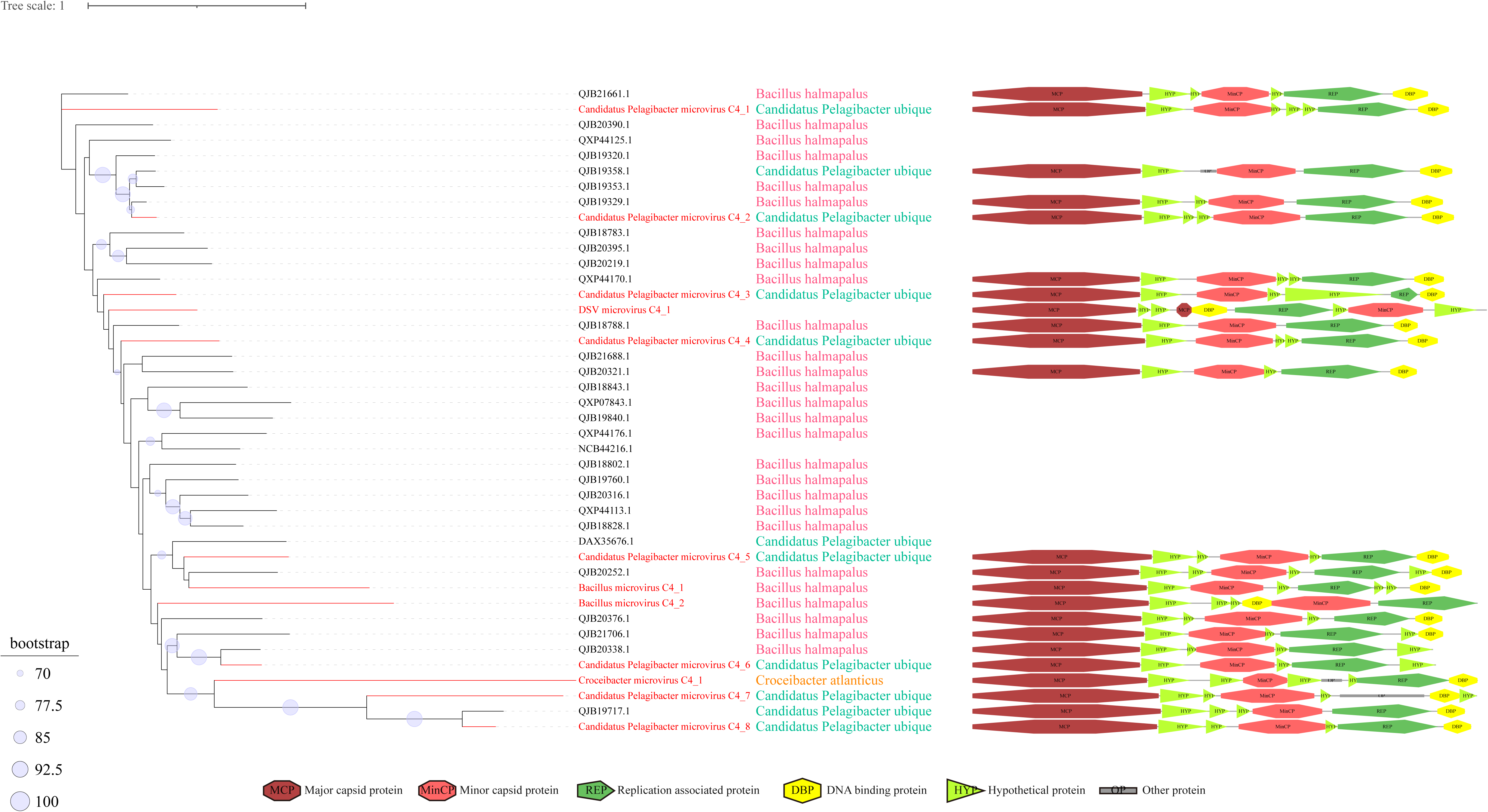

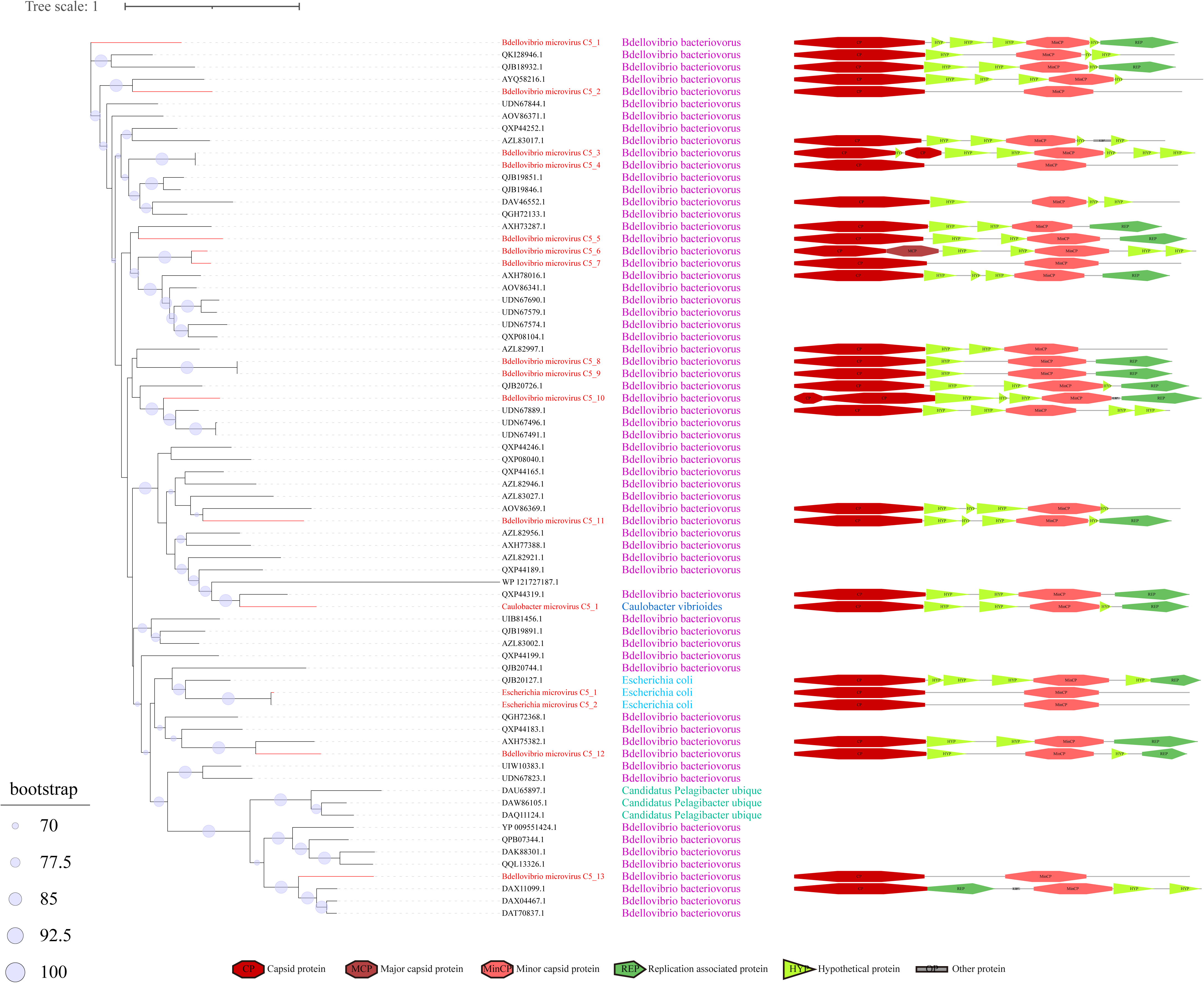

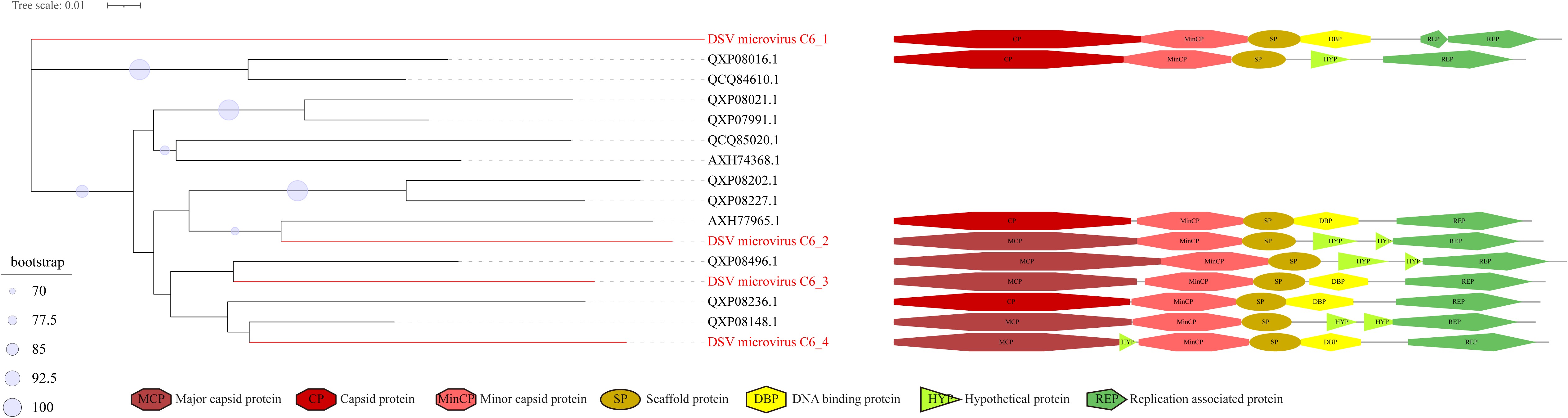

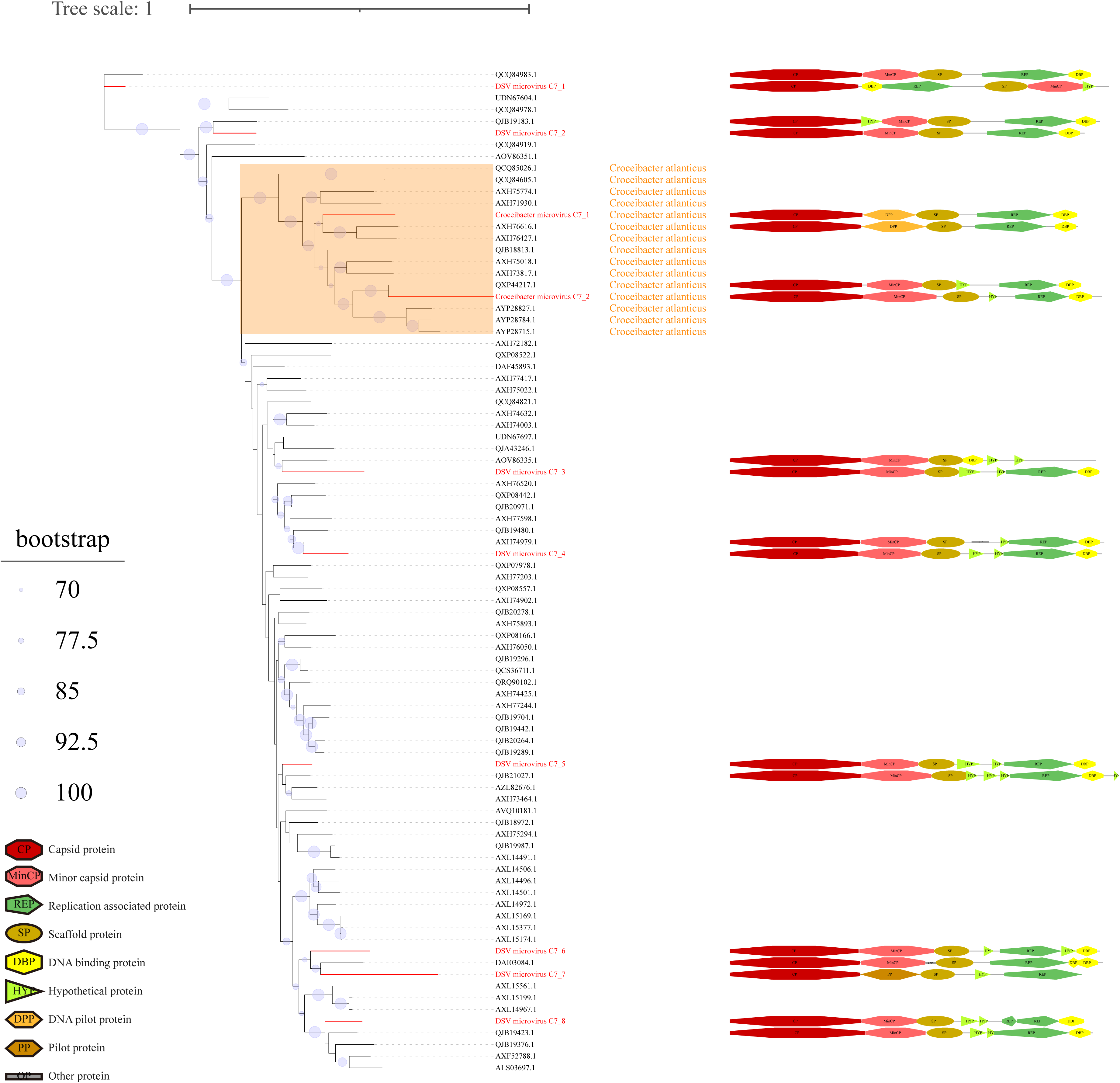

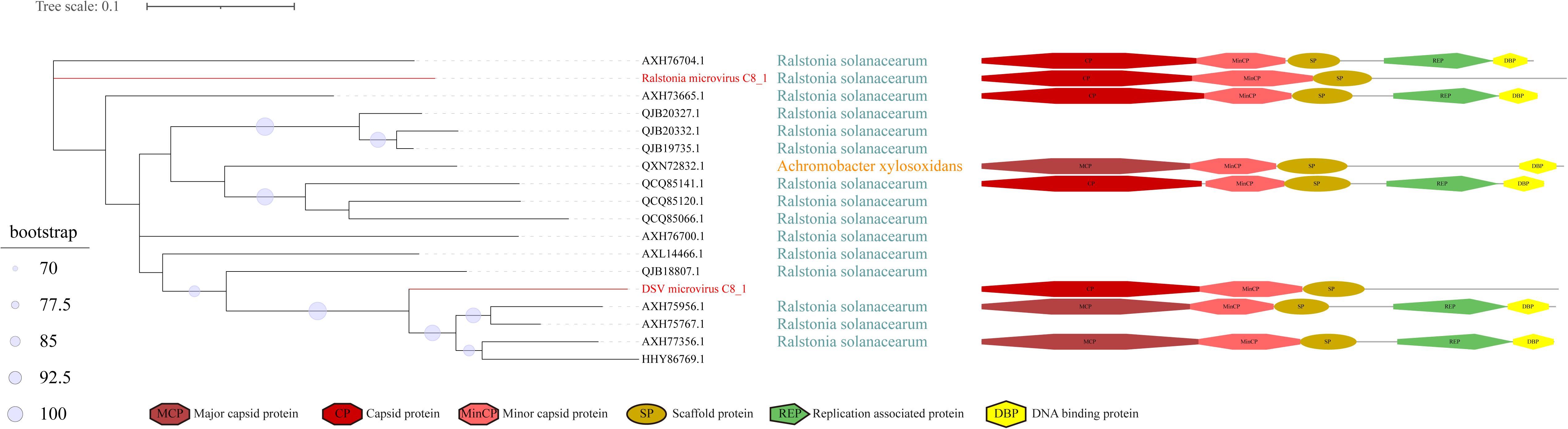

